# Prior aerosol infection with lineage A SARS-CoV-2 variant protects hamsters from disease, but not reinfection with B.1.351 SARS-CoV-2 variant

**DOI:** 10.1101/2021.05.05.442780

**Authors:** Claude Kwe Yinda, Julia R. Port, Trenton Bushmaker, Robert J. Fischer, Jonathan E. Schulz, Myndi G. Holbrook, Carl Shaia, Emmie de Wit, Neeltje van Doremalen, Vincent J. Munster

## Abstract

The circulation of SARS-CoV-2 has resulted in the emergence of variants of concern (VOCs). It is currently unclear whether previous infection with SARS-CoV-2 provides protection against reinfection with VOCs. Here, we show that low dose aerosol exposure to hCoV-19/human/USA/WA-CDC-WA1/2020 (WA1, lineage A), resulted in a productive mild infection. In contrast, low dose of SARS-CoV-2 via fomites did not result in productive infection in the majority of exposed hamsters and these animals remained non-seroconverted. After recovery, hamsters were re-exposed to hCoV-19/South African/KRISP-K005325/2020 (VOC B.1.351) via an intranasal challenge. Seroconverted rechallenged animals did not lose weight and shed virus for 3 days. They had little infectious virus and no pathology in the lungs. In contrast, shedding, weight loss and extensive pulmonary pathology caused by B.1.351 replication was observed in the non-seroconverted animals. The rechallenged seroconverted animals did not transmit virus to naïve sentinels via direct contact transmission, in contrast to the non-seroconverted animals. Reinfection with B.1.351 triggered an anamnestic response that boosted not only neutralizing titers against lineage A, but also titers against B.1.351. Our results confirm that aerosol exposure is a more efficient infection route than fomite exposure. Furthermore, initial infection with SARS-CoV-2 lineage A does not prevent heterologous reinfection with B.1.351 but prevents disease and onward transmission. These data suggest that previous SARS-CoV-2 exposure induces partial protective immunity. The reinfection generated a broadly neutralizing humoral response capable of effectively neutralizing B.1.351 while maintaining its ability to neutralize the virus to which the initial response was directed against.

## Introduction

SARS-CoV-2 transmission appears largely driven by direct contact and aerosol exposures. In experimental SARS-CoV-2 animal models, the exposure route and dose are linked to disease severity [1,2]. The evolution of SARS-CoV-2 has led to the emergence of several new variants. Within these novel emerging SARS-CoV-2 variants, there are variants of interest (VOIs) and variants of concern (VOCs). VOCs are defined by phenotypic changes including enhanced transmission, increased pathogenicity and decreased efficacy of prophylactic and therapeutic countermeasures [3,4]. Some VOCs display specific mutations in the spike protein that reduce binding affinity of neutralizing antibodies [5-8]. The relative protection induced by a previous SARS-CoV-2 infection against homologous or heterologous challenge is not fully understood. VOC B.1.351 (20H/501Y.V2) was first detected in South Africa and has been shown to exhibit reduced susceptibility to neutralization by sera from COVID-19 patients and vaccinated individuals [9,10]. Compared to the original Wuhan strain (lineage A), B.1.351 has nine amino acid substitutions and one deletion in the spike protein, including three changes in the receptor binding domain (RBD); K417N/T, E484K and N501Y [11].

We previously demonstrated the susceptibility of the Syrian hamster to SARS-CoV-2 aerosol and fomite exposure [2]. Here, we show that hamsters can productively be infected with a very low dose of aerosolized SARS-CoV-2, whereas a comparable low dose of fomite exposure did not result in infection. We then rechallenged the animals with B.1.351 to examine the potential for SARS-CoV-2 reinfection and onward transmission.

## Materials and Methods

### Ethics statement

Approval of animal experiments was obtained from the Institutional Animal Care and Use Committee of the Rocky Mountain Laboratories. Performance of experiments was done following the guidelines and basic principles in the United States Public Health Service Policy on Humane Care and Use of Laboratory Animals and the Guide for the Care and Use of Laboratory Animals. Work with infectious SARS-CoV-2 strains under BSL3 conditions was approved by the Institutional Biosafety Committee (IBC). Inactivation and removal of samples from high containment was performed per IBC-approved standard operating procedures.

### Virus strains and cell lines

SARS-CoV-2 strain hCoV-19/human/USA/WA-CDC-WA1/2020 (WA1, lineage A) was provided by CDC, Atlanta, USA while SARS-CoV-2 variant hCoV-19/South Africa/KRISP-K005325/2020 (B.1.351) was obtained from Dr. Tulio de Oliveira and Dr. Alex Sigal at the Nelson R. Mandela School of Medicine, UKZN. Both variants were propagated in VeroE6 cells in DMEM supplemented with 2% fetal bovine serum (FBS), 2 mM L-glutamine, 100 U/ml penicillin and 100 μg/ml streptomycin (DMEM2). VeroE6 cells were maintained in DMEM supplemented with 10% FBS, 2 mM L-glutamine, 100 U/ml penicillin and 100 μg/ml streptomycin. VeroE6 cells were provided by Dr. Ralph Baric. No mycoplasma was detected. Both strains were deep-sequenced, and no fixed mutations were found in nCoV-WA1-2020; the viral stock sequence was identical to the original patient sequence MN985325. However, for hCoV-19/South Africa/KRISP-K005325/2020 the viral stock contained the following non-fixed substitutions in the spike protein: Q677H (present at 88%) and R682W (present at 81%).

### Animal exposure and inoculation

Four to six-week-old female Syrian hamsters (ENVIGO) were exposed either via the intranasal, aerosol or fomite route as previously described by Port *et al*. 2020 [2]. For aerosol exposure non-anesthetized hamsters (4 per group) were placed into a stainless-steel wire mesh cage and subjected to 2.5×10^1^ and 1.1×10^2^ TCID_50_ SARS-CoV-2 aerosol particles, respectively, for 10 min using a 3-jet collision nebulizer (Biaera technologies, USA). Particles ranged from 1-5 µm in size. Dosage was confirmed by titration of the inoculum. Fomite exposure was conducted by exposing individually housed animals to a polypropylene dish containing 40 µl of 10^1^, 10^2^, and 10^3^ TCID_50_ SARS-CoV-2 per hamster (4 hamsters per group). At 14 days post inoculation (DPI) with aerosolized virus or fomites, blood was obtained by retro-orbitally bleeding and seroconversion was confirmed. At 21 DPI, 9 seroconverted and 2 non-seroconverted hamsters were intranasally rechallenged with 40 µl sterile DMEM containing 8×10^4^ TCID_50_ SARS-CoV-2 variant CoV-19/South African/KRISP-K005325/2020 (rechallenge). In all experiments, hamsters were individually housed. Weights were collected and oropharyngeal swabs were taken in 1 ml DMEM with 200 U/ml penicillin and 200 µg/ml streptomycin. At 5 days post rechallenge (DPR), 5 hamsters were euthanized, and lungs and serum collected. The remaining 6 animals were euthanized at 14 DPR. To assess transmission, all rechallenged animals were co-housed with a naïve sentinel (1:1 ratio) in a new cage for 24 h at 2 DPR. Sentinels were swabbed for three days.

### Viral RNA detection

Swabs from hamsters were collected as described above. RNA was extracted (140 µl) using the QIAamp Viral RNA Kit (Qiagen) using QIAcube HT automated system (Qiagen) according to the manufacturer’s instructions. Sub-genomic (sg) viral RNA was detected by qRT-PCR [12]. Five μl RNA (150 µl elution volume) was tested with TaqMan™ Fast Virus One-Step Master Mix (Applied Biosystems) using QuantStudio 6 Flex Real-Time PCR System (Applied Biosystems) according to the manufacturer’s instructions. Copy numbers/ml was calculated based on RNA standards.

### Virus titration

Viable virus in lungs or swabs was determined as previously described [13]. Briefly, tissues were weighted, then homogenized in 1 ml of DMEM2. Virus titrations were performed by end-point titration in VeroE6 cells, inoculated with tenfold serial dilutions of hamster swabs or tissue homogenates in 96-well plates. For swabs, plates were spun down for 1 h at 1000 rpm before incubation. Cytopathic effect was scored at day 6. TCID_50_ was calculated by the method of Spearman-Karber and, if required, adjusted for tissue weight.

### Histopathology

Necropsies and tissue sampling were performed according to IBC-approved protocols. Tissues were fixed for a minimum of 7 days in 10% neutral buffered formalin with 2 changes. Tissues were placed in cassettes and processed with a Sakura VIP-6 Tissue Tek, on a 12-hour automated schedule, using a graded series of ethanol, xylene, and ParaPlast Extra. Prior to staining, embedded tissues were sectioned at 5 µm and dried overnight at 42°C. Using GenScript U864YFA140-4/CB2093 NP-1 (1:1000) specific anti-CoV immunoreactivity was detected using the Vector Laboratories ImPress VR anti-rabbit IgG polymer (# MP-6401) as secondary antibody. The tissues were then processed using the Discovery Ultra automated processor (Ventana Medical Systems) with a ChromoMap DAB kit Roche Tissue Diagnostics (#760-159).

### Serology - ELISA

Serum samples were inactivated with γ-irradiation (2 mRad). Maxisorp plates (Nunc) were coated with 50 ng spike protein per well and incubated overnight at 4°C. After blocking with casein in phosphate buffered saline (PBS) (ThermoFisher) for 1 h at room temperature (RT), serially diluted 2-fold serum samples (duplicate, in casein) were incubated for 1 h at RT. Spike-specific antibodies were detected with goat anti-hamster IgG Fc (horseradish peroxidase (HRP)-conjugated, Abcam) for 1 h at RT and visualized with KPL TMB 2-component peroxidase substrate kit (SeraCare, 5120-0047). The reaction was stopped with KPL stop solution (Seracare) and read at 450 nm. Plates were washed 3x with PBS-T (0.1% Tween) in between steps. The threshold for positivity was calculated as the average plus 3x the standard deviation of negative control hamster sera.

### Serology - Virus neutralization

Heat-inactivated γ-irradiated sera were two-fold serially diluted in DMEM2. 100 TCID_50_ of SARS-CoV-2 strain nCoV-WA1-2020 or hCoV-19/South African/KRISP-K005325/2020 was added. After 1 h of incubation at 37°C and 5% CO_2_, the virus:serum mixture was added to VeroE6 cells. CPE was scored after 5 days at 37°C and 5% CO_2_. The virus neutralization titer was expressed as the reciprocal value of the highest dilution of the serum which still inhibited virus replication.

## Results

### Low-dose aerosol exposure is highly efficient and displays a mild disease phenotype in hamsters

We previously demonstrated that hamsters become uniformly infected after an experimental aerosol or fomite exposure with a relatively high dose of 8×10^4^ TCID_50_ SARS-CoV-2 [2]. To increase our understanding of the relationship between exposure route and infectious dose we exposed Syrian hamsters to SARS-CoV-2 administered via low dose aerosols or fomites. For the aerosol inoculation route, two groups (N = 4) of hamsters were exposed to 1.1×10^2^ or 2.5×10^1^ TCID_50_ of SARS-CoV-2 strain nCoV-WA1-2020 (WA1, lineage A; indicated as Aerosol 10^2^ and 10^1^). For the fomite inoculation route, three groups (N = 4) of hamsters were exposed to 10^3^, 10^2^ and 10^1^ TCID_50_ of SARS-CoV-2 WA1 (indicated as Fomite 10^3^, 10^2^ and 10^1^). No significant median weight loss was observed following inoculation for any of the groups (**Fig 1 a**). To assess productive infection, we measured subgenomic RNA (sgRNA) in oropharyngeal swabs at 1, 2, 3, 5, 7, and 9 DPI. In the Aerosol 10^2^ and 10^1^ groups, respiratory shedding started at 1 DPI and lasted 9 days on average. In contrast, no prolonged shedding was observed in the Fomite 10^1^ and 10^3^ groups, and only one animal in the Fomite 10^2^ group demonstrated prolonged shedding until 9 DPI (**Fig 1b**). To compare the magnitude of shedding between the two aerosol groups, we performed an area under the curve (AUC) analysis. No significant difference between the two groups was observed (median Aerosol 10^2^ = 46.16 and Aerosol 10^1^ = 60.57 AUC log_10_ copies/exp, N = 4, Mann-Whitney test, p = 0.0571) (**Fig 1 c**). All aerosol-exposed hamsters seroconverted by 14 DPI, as measured by anti-spike ELISA. In contrast, seroconversion was only detected in the single animal which demonstrated prolonged shedding in the Fomite 10^2^ group. The magnitude of the humoral response was higher in Aerosol 10^2^ animals than in the Aerosol 10^1^ animals (**Fig 1 d**, median Aerosol 10^2^ = 409,600 and Aerosol 10^1^ = 51,200 ELISA titer, N = 4, Mann Whitney test, p = 0.0286). This study confirms that aerosol exposure is a highly efficient infection route in hamsters, whereas fomite exposure is less so.

**Figure 1.**
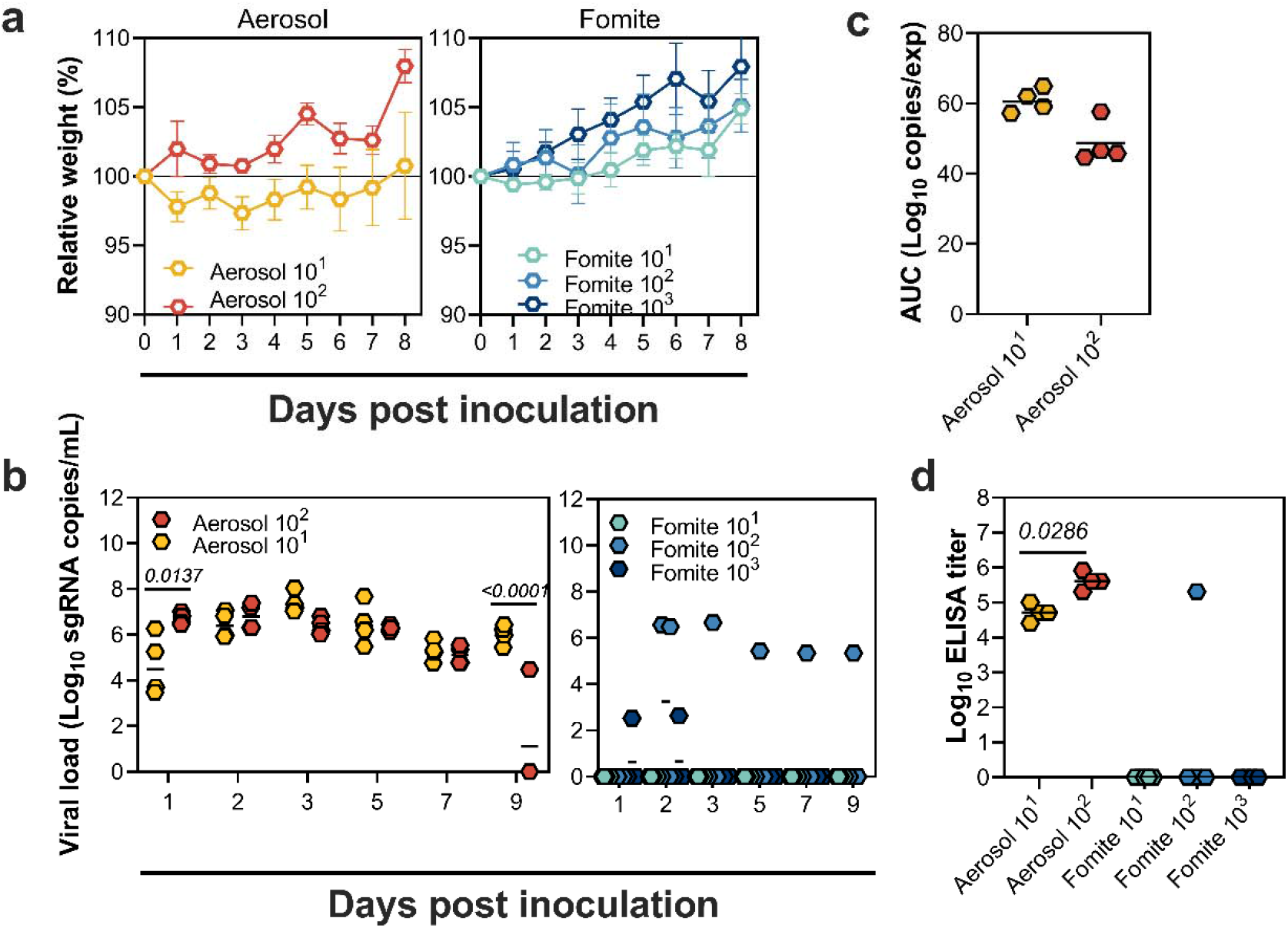
Infection efficiency of low-dose SARS-CoV-2 aerosol and fomite exposure. Relative weight in hamsters after SARS-CoV-2 inoculation via the aerosol route with 1.1×10^2^ or 2.5×10^1^ TCID_50_ (indicated as Aerosol 10^2^ and 10^1^) or fomite route with 10^3^, 10^2^ and 10^1^ TCID_50_ (indicated as Fomite 10^3^, 10^2^ and 10^1^) of hCoV-19/human/USA/WA-CDC-WA1/2020 (WA1, lineage A), (N = 4 per group). Symbols show mean and error bars show standard error of the mean (SEM). b. Viral load in oropharyngeal swabs of individual animals. c. Area under the curve (AUC) analysis of sgRNA detected in oropharyngeal swabs throughout the experiment. The lines represent the median. d. Binding antibodies against spike protein of SARS-CoV-2 in serum obtained at 14 days post inoculation.

### Previous low-dose aerosol exposure with SARS-CoV-2 lineage A protects hamsters from disease after high-dose B.1.351 challenge

Next, we investigated whether reinfection with VOC B.1.351 would occur after a prior low-dose aerosol or fomite exposure with lineage A. At 21 DPI, nine seroconverted animals (four of each aerosol exposed group and one Fomite 10^2^ animal), and two non-seroconverted animals (Fomite 10^1^ group) were each intranasally inoculated with 8×10^4^ TCID_50_ VOC B.1.351. Seroconverted animals did not lose weight after rechallenge. In contrast, the two naive animals lost weight at 5 DPR (**Fig 2 a**). Compared to the non-seroconverted animals, the seroconverted animals shed relatively low amounts of sgRNA and for a shorter duration. The magnitude of shedding was higher in non-seroconverted animals (median seroconverted = 16.62 (N = 9) and non-seroconverted = 30.42 AUC log_10_ copies/exp (N = 2)) (**Fig 2 b/c**). At 5 DPR, three seroconverted animals and the two non-seroconverted animals were selected for necropsy. Lung:body weight ratio was increased in non-seroconverted animals compared to seroconverted animals (median seroconverted = 0.7256 mg/g (N = 3) and non-seroconverted = 1.057 mg/g (N = 2)) (**Fig 2 d**).

**Figure 2.**
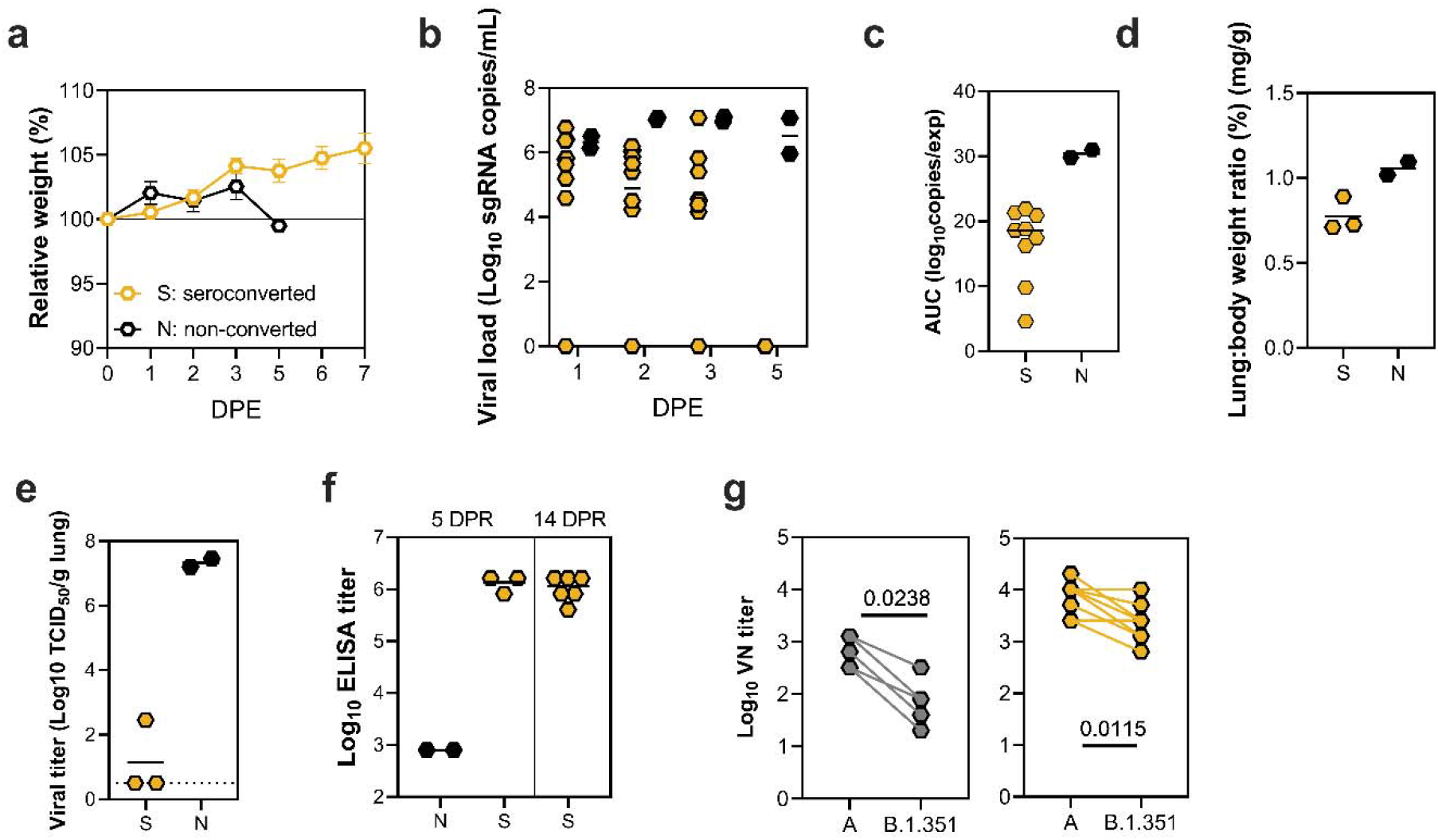
Heterologous B.1.351 challenge following SARS-CoV-2 lineage A exposure. a. Relative weight in hamsters after 8×10^4^ TCID_50_ VOC B.1.351 intranasal rechallenge over time (DPR= day post rechallenge) for the seroconverted animals (N = 9) and non-seroconverted animals (N = 2). Symbols show mean and error bars show standard error of the mean (SEM). b. Viral load in oropharyngeal swabs from individual rechallenged animals. Line represents median. c. Area under the curve (AUC) analysis of shedding as measured by viral load in swabs. Line represents median. d. Lung:body weight ratio (mg/g) of hamsters euthanized at 5 DPR. Line represents median. e. Infectious virus titer in lung tissue obtained at 5 DPR. Line represents median, dotted line represents limit of detection. f. Binding antibodies against spike protein of SARS-CoV-2 in serum obtained 14 DPR. g. Left panel: Virus neutralizing antibody titers against lineage A and VOC B.1.351. Right panel: Virus neutralizing antibody titers against lineage A and VOC B.1.351 in serum obtained from seroconverted animals after VOC B.1.351 rechallenge.

Up to 10^7^ TCID_50_ SARS-CoV-2/g tissue was detected in the lungs of non-seroconverted animals while only one of the seroconverted animals was positive for infectious virus with a titer of 10^2.5^ TCID_50_/g (**Fig 2 e**). Spike protein amino acid substitutions Q677H and R682W were completely cleared from the virus population and no longer detected in lung tissue of virus positive animals. Upon histopathological analysis lung tissue from seroconverted animals was normal (**3 a, b, e, f)** while the lungs of the 2 non-seroconverted animals demonstrated alveolar inflammation consisting of macrophages and neutrophils, as well as variable amounts of hemorrhage, fibrin and edema (**Fig 3 c, d, g, h**) consistent with a lower respiratory tract infection with SARS-CoV-2. Immunohistochemistry using a monoclonal antibody against SARS-CoV-2 demonstrated viral antigen in bronchial and bronchiolar epithelium, type I and II pneumocytes as well as pulmonary macrophages within the non-seroconverted animals (**Fig 3 k, l)**, but not in the seroconverted animals (**Fig 3 i, j)**.

**Figure 3.**
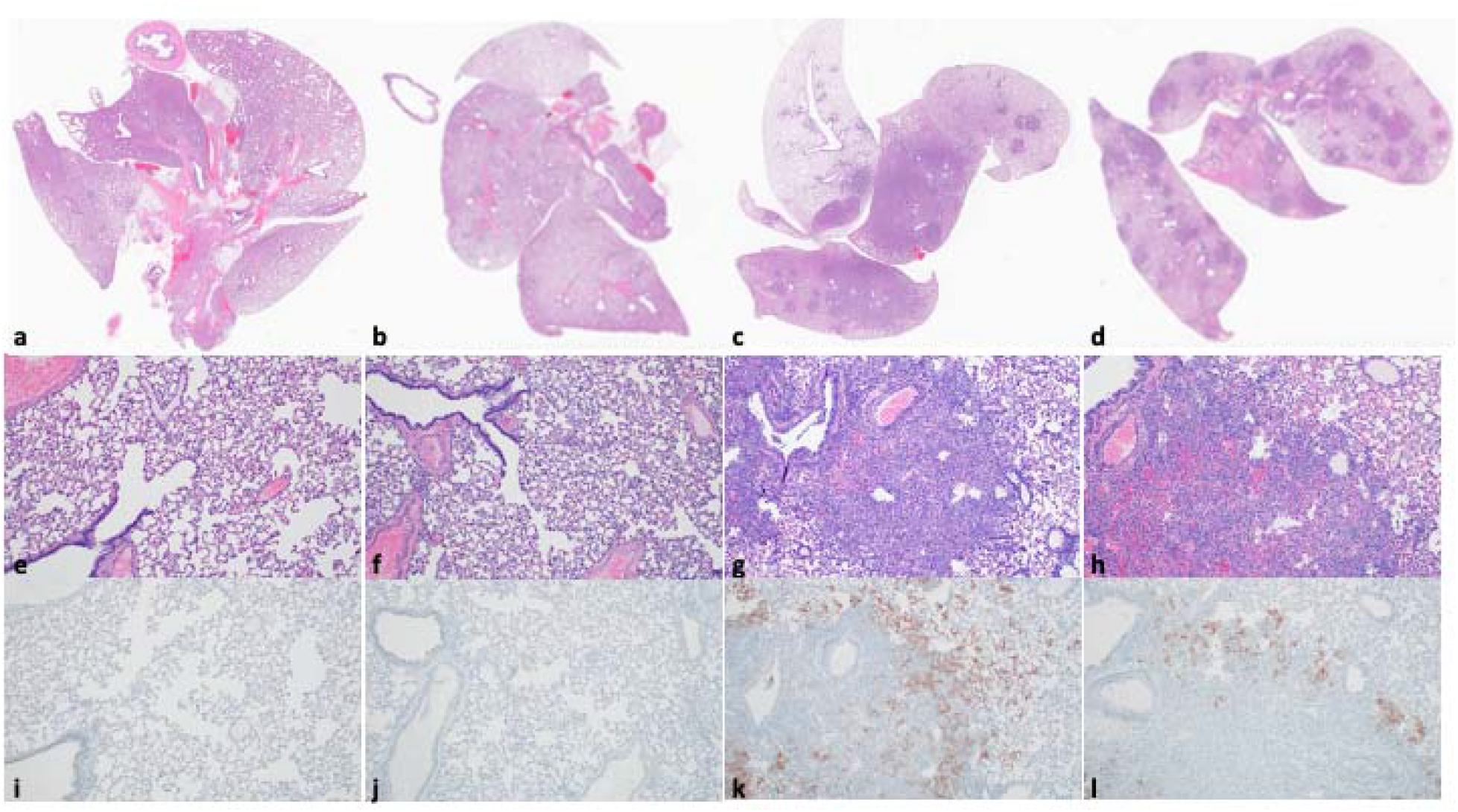
Lung pathology after heterologous B.1.351 rechallenge. Comparison of SARS-CoV-2 pathology for seroconverted and non-seroconverted hamsters at 5 days post rechallenge (DPR). a-d. HE 1x. Lungs of seroconverted hamsters (a, b) and non-seroconverted hamsters (c, d), lung tissue from seroconverted animals was normal whereas the non-seroconverted animals displayed pathology consistent of lower respiratory tract infection with SARS-CoV-2. e-h. HE 100x. Normal lungs from seroconverted hamsters (e, f) and lungs from non-seroconverted hamsters demonstrated alveolar inflammation consisting of macrophages and neutrophils, as well as variable amounts of hemorrhage, fibrin and edema (g, h). i-l. IHC staining against N protein SARS-CoV-2 (SARS-CoV-2 antigen is visible as red-brown staining), 100x. SARS-CoV-2 antigen is absent from lungs of the seroconverted hamsters (i, j) whereas lungs from non-seroconverted hamsters demonstrated viral antigen in bronchial and bronchiolar epithelium, type I and II pneumocytes as well as pulmonary macrophages (k, l).

Rechallenge with VOC B.1.351 led to an increase in the humoral response. ELISAs conducted at 5 DPR showed seroconverted rechallenged animals had increased levels of antibody titers against lineage A SARS-CoV-2 spike protein (median = 1,638,400 ELISA titer, N = 3) compared to the minimal levels of antibodies in the two non-seroconverted animals (median = 800, N = 2) (**Fig 2 f**). Next, we investigated neutralizing antibody titers in the serum of initial and rechallenged animals against SARS-CoV-2 lineage A and B.1.351 variants. Neutralization of B.1.351 was significantly reduced compared to neutralization of SARS-CoV-2 lineage A (median = 80/640 VN titer, median 16-fold reduction, N = 5, Mann-Whitney test, p = 0.0238, **Fig 2 g**). The rechallenge with VOC B.1.351 resulted in an increased neutralizing titer against both B.1.351 (median = 2,560 VN titer, N = 9) and lineage A (median = 10,240 VN titer, N = 9). However, neutralization of B.1.351 was still significantly reduced compared to neutralization of SARS-CoV-2 lineage A (median 4-fold reduction, N = 9, Mann-Whitney test, p = 0.0115).

These data suggest that prior exposure does not protect hamsters from reinfection with an antigenically different SARS-CoV-2 virus. However, the humoral immune response generated by a low dose primary infection provides protection against disease and lower respiratory tract replication with an antigenically different SARS-CoV-2 virus and the replication remains confined to the upper respiratory tract.

### Previous low-dose aerosol exposure with SARS-CoV-2 lineage A protects hamsters from transmitting VOC B.1.351 to naïve animals

We next observed the ability of rechallenged hamsters to transmit VOC B.1.351 to naïve hamsters. Eight seroconverted and two non-seroconverted rechallenged hamsters (N = 10) were co-housed with naive sentinels at 2 DPR for 24 h. We assessed transmission by taking oropharyngeal swabs of the sentinels at day 1, 2 and 3 post exposure (DPE) and analysing them for the presence of sgRNA. In the two sentinels co-housed with a non-seroconverted rechallenged hamster, we observed shedding of sgRNA from the sentinel hamsters (**Fig 4 a**). In contrast, in swabs obtained from sentinels co-housed with seroconverted rechallenged hamsters, no SARS-CoV-2 RNA was detected at any timepoint suggesting that no transmission occurred. The efficiency of transmission was also evaluated by serology at 14 DPE. Only the two sentinels co-housed with a non-seroconverted rechallenged hamster had seroconverted, as measured by anti-spike ELISA (**Table 1**). We thus conclude that none of the seroconverted animals transmitted B.1.351 to sentinels, whereas both non-seroconverted animals transmitted virus to sentinels.

**Table 1:** Presence of SARS-CoV-2 spike IgG antibodies in sentinels co-housed with B.1.351 rechallenged hamsters in serum obtained 14 days post contact. negative (-): optical density (at 450 nm) < 0.124, positive (+) optical density (at 450 nm) ≥ 0.124.

| Donor | ID | Seroconversion status |
| --- | --- | --- |
| Seroconverted | Sentinel 1 | (-) |
|  | Sentinel 2 | (-) |
|  | Sentinel 3 | (-) |
|  | Sentinel 4 | (-) |
|  | Sentinel 5 | (-) |
|  | Sentinel 6 | (-) |
|  | Sentinel 7 | (-) |
|  | Sentinel 8 | (-) |
| Non-seroconverted | Sentinel 9 | (+) |
|  | Sentinel 10 | (+) |

**Figure 4.**
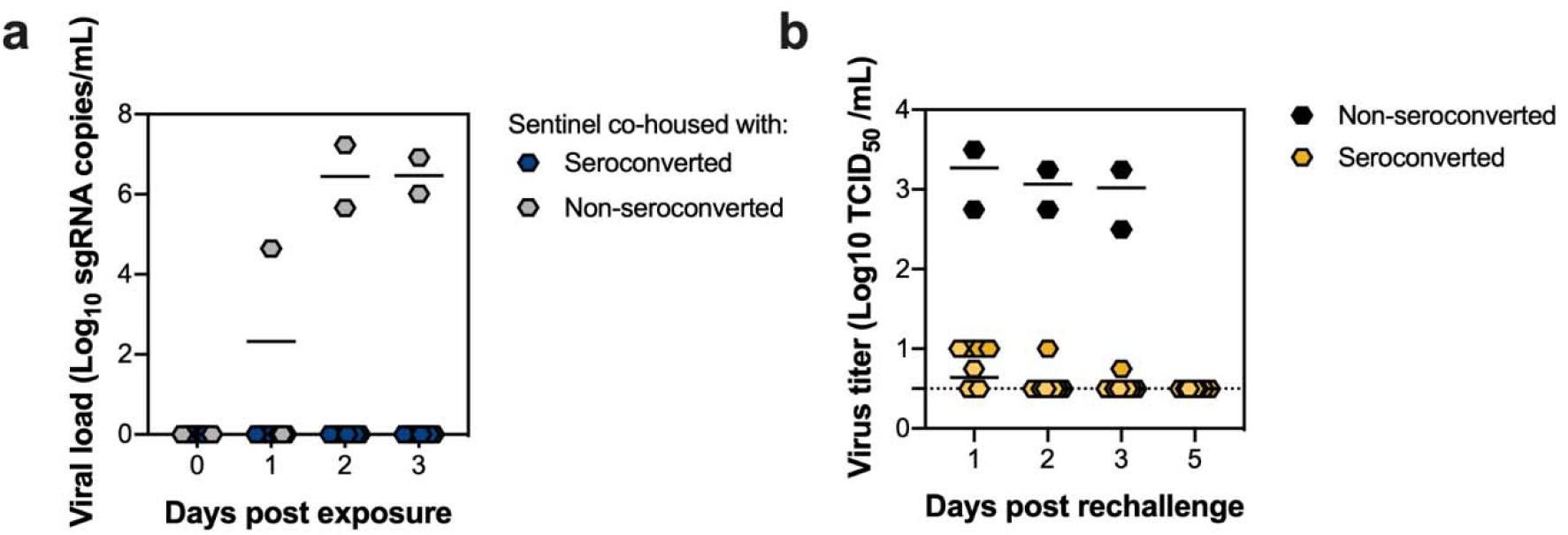
Hamsters rechallenged with VOC B.1.351 do not transmit SARS-CoV-2 to naïve sentinels. VOC B.1.351 rechallenged animals were co-housed with a naïve sentinel (1:1 ratio) in a new cage for 24 h at 2 DPR. a. Viral load of individual exposed sentinel animals measured by sgRNA in oropharyngeal swabs days post exposure (DPE). b. Infectious virus titer of individual animals in oropharyngeal swabs from rechallenged animals. Dotted line = limit of detection.

To understand why transmission did not occur for sentinels co-housed with seroconverted hamsters, we determined the level of infectious virus in oropharyngeal swabs obtained from donor animals. Infectious virus could be detected in swabs from both non-seroconverted animals at 2 DPR (median = 1170 TCID_50_/ml, N = 2), while infectious virus was detected in only one of the seroconverted animals and at a very low level (10 TCID_50_/ml) **(Fig 4 b)**. This indicates that previous challenge with lineage A by low-dose aerosol and fomite leads to rapid upper respiratory tract clearance of infectious virus after rechallenge and subsequently limits transmission potential.

## Discussion

We investigated SARS-CoV-2 infection of Syrian hamsters after low dose exposure to aerosols or fomites. We demonstrated for the first time that aerosol exposure of only 2.5×10^1^ TCID_50_ SARS-CoV-2 was sufficient to cause productive infection in Syrian hamsters. In addition, we confirmed the inefficiency of infection of hamsters after fomite exposure. These results are in line with previous experimental studies where infection via airborne SARS-CoV-2 or influenza A virus was more efficient than infection through contaminated environmental surfaces [14,15].

In humans, the relative importance of each transmission route remains unclear, but the general consensus is that the principal mode of infection is via respiratory droplets and that the relative risk of fomite transmission of SARS-CoV-2 is considered low compared with direct contact, droplet transmission, or airborne transmission [16-20]. Our experimental data is in line with human epidemiological case studies which show that in closed spaces airborne transmission is highly efficient [21-23].

The emergence of VOC B.1.351 in South Africa has raised concerns with regards to its reinfection potential due to antigenic drift. VOC B.1.351 shows significant reduction of neutralizing titers in human convalescent serum [24-30]. The degree of protective immunity by previous SARS-CoV-2 infection in humans is currently unknown, but several reports suggest that reinfection does occasionally occur [31-33]. In the Syrian hamster model, homologous challenge resulted in reduced virus shedding and reduced replication in the lungs [34,35]. In addition, homologous rechallenged hamsters were unable to transmit to naïve animals [35]. Our reinfection data confirm the prior observations and also show that heterologous reinfection with VOC B.1.351 results in limited replication in the lungs, absence of pathology and prevention of onwards transmission in a direct contact transmission model.

Protection against VOC B.1.351 rechallenge was demonstrated in the Syrian hamster model after prior intranasal infection with a lineage A virus [36]. However, this study relied on a high-dose intranasal infection with lineage A. Our study evaluated reinfection potential of VOC B.1.351 after low-dosage fomite or aerosol exposure with lineage A. The B.1.351 virus stock used to challenge hamsters contained non-fixed amino acid variation at position Q677H and R682W (88% and 81%, respectively). These major variant amino acid SNPs were absent on 5 DPR, suggesting that they are rapidly selected against in the SARS-CoV-2 hamster model. Nonetheless, the substitutions which are thought to be important in immune evasion, such as E484K, are still present in the virus stock. Furthermore, efficient upper and lower respiratory tract replication and lung pathology was observed in infected hamsters. We thus do not believe that the presence of these additional amino acid substitutions, present as a quasispecies, affect the data interpretation.

Our results provide more experimental support to the consensus that aerosol exposure is a more efficient infection route than fomite exposure [37]. Whereas initial infection with SARS-CoV-2 lineage A did not directly prevent heterologous reinfection with VOC B.1.351, it did reduce shedding, and did prevent disease and lower respiratory tract replication and onward transmission. Several studies have reported reduced neutralization of B.1.351 by convalescent plasma and vaccine-induced neutralizing antibodies [38-41] and here we confirm this observation with hamster sera with a 16-fold drop in neutralizing titer between a lineage A virus and the B.1.351. Reinfection with B.1.351 triggered an anamnestic response that boosted not only the neutralizing titer against lineage A, but also against B.1.351, and the difference between neutralizing capacity was reduced from 16-fold to 4-fold. These data suggest that reinfection with B.1.351 not only boosts the existing antibody repertoire, but in addition generates a broadly neutralizing humoral response. This humoral response is capable of effectively neutralizing B.1.351 while maintaining its ability to neutralize the virus to which the initial response was generated against. Finally, the induction of partial protective immunity by previous SARS-CoV-2 exposure might reduce the risk for lower respiratory tract COVID-19 disease and onward transmission in humans.

## Acknowledgements

We would like to thank Brandi Williamson, Craig Martens, Kent Barbian, Stacey Ricklefs, Sarah Anzick and the animal care takers for their excellent assistance during the study and Sujatha Rashid, Ranjan Mukul, Kimberly Stemple for critical help with obtaining the SARS-CoV-2 isolates in this study. The following reagent was obtained through BEI Resources, NIAID, NIH: SARS-Related Coronavirus 2, Isolate hCoV-19/South Africa/KRISP-K005325/2020, NR-54009, contributed by Alex Sigal and Tulio de Oliveira. This research was supported by the Intramural Research Program of the National Institute of Allergy and Infectious Diseases (NIAID), National Institutes of Health (NIH).

